# Design, expression, purification and characterization of a YFP-tagged 2019-nCoV spike receptor-binding domain construct

**DOI:** 10.1101/2020.09.29.318196

**Authors:** Tobias Bierig, Gabriella Collu, Alain Blanc, Emiliya Poghosyan, Roger. M. Benoit

## Abstract

2019-nCoV is the causative agent of the serious, still ongoing, worldwide COVID-19 pandemic. High quality recombinant virus proteins are required for research related to the development of vaccines and improved assays, and to the general understanding of virus action. The receptor-binding domain (RBD) of the 2019-nCoV spike (S) protein contains disulfide bonds and N-linked glycosylations, therefore, it is typically produced by secretion. Here, we describe a construct and protocol for the expression and purification of yellow fluorescent protein (YFP) labeled 2019-nCoV spike RBD. The fusion protein, in the vector pcDNA 4/TO, comprises an N-terminal interferon alpha 2 (IFNα2) signal peptide, an eYFP, a FLAG-tag, a human rhinovirus 3C protease cleavage site, the RBD of the 2019-nCoV spike protein and a C-terminal 8x His-tag. We stably transfected HEK 293 cells. Following expansion of the cells, the fusion protein was secreted from adherent cells into serum-free medium. Ni-NTA IMAC purification resulted in very high protein purity, based on analysis by SDS-PAGE. The fusion protein was soluble and monodisperse, as confirmed by size-exclusion chromatography (SEC) and negative staining electron microscopy. Deglycosylation experiments confirmed the presence of N-linked glycosylations in the secreted protein. Complex formation with the peptidase domain of human angiotensin-converting enzyme 2 (ACE2), the receptor for the 2019-nCoV spike RBD, was confirmed by SEC, both for the YFP-fused spike RBD and for spike RBD alone, after removal of YFP by proteolytic cleavage. Possible applications for the fusion protein include binding studies on cells or *in vitro*, fluorescent labeling of potential virus-binding sites on cells, the use as an antigen for immunization studies or as a tool for the development of novel virus- or antibody-detection assays.

## Introduction

The membrane-anchored, trimeric spike (S) glycoproteins are the most prominent protrusions on the surface of 2019-nCoV. CoV spike proteins typically comprise two subunits. The S1 subunit is responsible for receptor binding and the S2 subunit is involved in fusing the membranes of the virus and the host (Li 2016). The S1 subunit is composed of an N-terminal domain (S1-NTD) and a C-terminal domain (S1-CTD) (Li 2016). S1-CTD comprises two subdomains, one functioning as a core structure, the other one as a receptor-binding motif (Li 2016). The receptor-binding domain of 2019-nCoV binds human angiotensin-converting enzyme 2 (ACE2) with high affinity (Wrapp et al. 2020). The spike RBD is an important target for drug discovery research (Toelzer et al. 2020) and for the development of vaccines (Wrapp et al. 2020; Wang et al. 2020).

Within the S trimer, the receptor-binding domains (RBDs) can be in a down conformation or alternatively in an up conformation, the latter being the receptor-accessible state (Wrapp et al. 2020). Recent complex structures (Yan et al. 2020; Wang et al. 2020) confirmed that a single spike RBD, taken out of the trimeric context, is capable of binding its human receptor ACE2.

Here, we describe a construct and protocol for the production and purification of milligram amounts of N-terminally YFP-labeled spike RBD. The domain boundaries of our receptor-binding domain (RBD) construct are based on the construct used for the crystal structure by Wang et al. (Cell 2020, PDB entry 6LZG), comprising amino acids 319-527, which also includes the receptor-binding motif (amino acids 437-508, UniProtKB - P0DTC2).

Expression is performed by secretion into serum-free medium from adherent, stably transfected HEK293 cells. The protocol involves only standard cell culture techniques and equipment. Our experiments confirmed that the fusion protein (also after proteolytic removal of YFP) binds the human ACE2 peptidase domain.

## Materials and Methods

### Plasmids

The DNA coding for the IFNα2-eYFP-FLAGtag-PreScission_site-S_RBD-8xHis-tag-StopStop fusion protein was ordered from Genewiz, cloned into the HindIII and XbaI sites of pcDNA 4/TO (Invitrogen). The human ACE2 peptidase domain (amino acids 19 - 615) construct with an N-terminal interleukin-2 (IL-2) peptide and a C-terminal 8xHis-tag and two stop codons was ordered as a FragmentGENE from Genewiz and cloned into the KpnI and NotI sites of pcDNA 4/TO.

### Transfection

Adherent HEK293 cells were grown to ∼90% confluence in a 9 cm diameter cell culture dish at 37°C, 5% CO_2_. Just before transfection, the cells were washed with 10 ml PBS. 22 µg plasmid DNA and 50 µg of 25 kDa, linear polyethylenimine (PEI) were mixed in a sterile 15 ml Falcon tube and incubated at room temperature for 10 minutes with occasional gentle mixing. Next, 5 ml of Dulbecco’s Modified Eagle Medium (DMEM), with high glucose and L-glutamine (Bioconcept), without FBS, were added to the DNA-PEI mixture, followed by another 10 minutes incubation at room temperature with occasional mixing. Thereafter, the PBS was removed from the cells and the transfection mixture was added onto the cells and distributed well. After incubation at 37°C, 5% CO_2_ for six hours, 10 ml of DMEM high glucose supplemented with 1% FBS were added, and the cells were incubated at 37°C, 5% CO_2_ overnight The next day, the cells were split 1:10 and grown in DMEM high glucose medium supplemented with 10% FBS.

### Selection of stable cell lines

After another overnight incubation, the medium was replaced by fresh medium of the same composition, and supplemented with Zeocin (InvivoGen) to a final concentration of 100 µg/ml and Penicillin-Streptomycin (PAN Biotech) to a final concentration of 100 U/ml. The selective medium was exchanged on Mondays, Wednesdays and Fridays until only Zeocin-resistant cells remained and the cells were confluent. Nine days after transfection, the cells were trypsinized and transferred to a 75 cm^2^ cell culture flask in 20 ml selective medium. Fourteen days after transfection, 3/5 of the cells in the 75 cm^2^ flask (∼25% confluent) were split into a new 75 cm^2^ flask for expression, while the other 2/5 were transferred to another flask as a backup and for freezing.

### Protein expression

Sixteen days after transfection, when the cells in the expression flask were ∼50% confluent, they were washed twice with PBS, and 15 ml of serum-free, selective expression medium (Opti-MEM I reduced serum medium, Gibco REF 11058-021) supplemented with 100 µg/ml Zeocin (InvivoGen cat no ant-zn) and 3 µg/ml tetracycline) were added. The selective expression medium was collected on Mondays, Wednesdays and Fridays and replaced with fresh medium of the same composition. Once in the serum-free medium, the cells reached confluence within 10 days. The supernatant medium collected from the confluent culture typically contained some detached cells, which were removed by centrifugation at room temperature, 1500 rcf for 10 minutes.

For upscaling, the cells were expanded into two cell culture flasks with 150 cm^2^ surface area each. The cells were grown to confluence and then washed twice in PBS before adding serum-free medium for expression.

### Protein purification

#### Ni-NTA IMAC

After removal of detached cells by centrifugation, the expression medium containing the secreted fusion protein was supplemented with 1/4 tablet of protease inhibitor (complete EDTA-free protease inhibitor cocktail tablets, Roche Diagnostic GmbH, 45148300) and transferred to a 15 ml Falcon tube containing 200 μl of washed, pre-equilibrated Ni-NTA agarose (Qiagen Cat. No. 30210). The solution was incubated at 4°C with occasional agitation until the next batch of secreted protein became available. Then, the Ni-NTA resin was collected by centrifugation at 1500 rcf, 10 minutes at 4°C. The supernatant was discarded and the fresh batch of clarified, protease inhibitor treated medium was added to the resin. This process was repeated until the Ni-NTA agarose clearly became yellow. The Ni-NTA resin was then collected by centrifugation (1500 rcf, 10 minutes, 4°C), the supernatant was discarded, and the resin was resuspended in wash buffer (50 mM Tris pH 8.0, 300 mM NaCl) and transferred to a column. The resin was washed six times with 1 ml of ice-cold wash buffer per wash, using gravity flow. The protein was eluted in three 300 μl steps in 50 mM Tris pH 8.0, 300 mM NaCl, 350 mM imidazole. For the upscaled purification, 1 ml of Ni-NTA agarose was used.

### SEC

The IMAC-purified proteins were centrifuged for 10 minutes at 17000 rcf, 4°C and run on a Superdex 200 Increase 10/300 GL column (Code 28-9909-44) in 50 mM Tris pH 8.0, 150 mM NaCl, at a flow rate of 0.4 ml/minute on an ÄKTA Ettan system at 4°C. For binding studies, separate proteins were centrifuged 10 minutes at 17000 rcf, 4°C. Equimolar amounts of the supernatants were then mixed and incubated on ice for 1 hour prior to the SEC run. For the complex containing the YFP-S_RBD fusion protein, 22 μg of SEC-purified YFP-S_RBD were mixed with an equimolar amount of IMAC-purified ACE2 peptidase domain, and the volume was adjusted to 500 μl using SEC buffer. For the complex containing cleaved S RBD without YFP, 34 μg of IMAC-purified S_RBD were mixed with an equimolar amount of IMAC-purified ACE2 peptidase domain, and the volume was adjusted to 500 μl using SEC buffer. After the incubation step, before the SEC run, the complexes were again centrifuged for 10 minutes at 17000 rcf, 4°C.

### Electron microscopy

#### Negative stain grid preparation

3.5 - 4 µl of purified protein (0.125 mg/mL) were first applied to a glow discharged, carbon coated grid (Plano, Germany), thereafter excess liquid was blotted away using filter paper and grids were stained with 1-2% uranyl acetate solution.

#### Cryo-EM grid preparation

The protein peak obtained from SEC was collected and concentrated to 0.6 mg/ml. Cryo-EM grids were prepared by applying 3.5 μl of protein to the glow-discharged Quantifoil R1.2/1.3 200-copper mesh grids from Electron Microscopy Science (Q2100-CR1.3). The grids were blotted for 3 s, plunge-frozen in liquid ethane using a Vitrobot Mark IV (Thermo Fischer Scientific), operated at 4°C and 100% humidity, and stored in liquid nitrogen until cryo-EM data collection.

#### EM data acquisition

Data acquisition was performed using a JEM2200FS transmission electron microscope (JEOL, Tokyo, Japan) equipped with an in-column energy filter and a field emission gun. Micrographs were recorded with K2/XP direct electron detector (Gatan, Ametek) and GMS3 software (Gatan, Ametek).

### Deglycosylation

For analysis by SDS-PAGE, 5 μl 1x PBS and 2 μl (1 U) PNGase F (from *Elizabethkingia meningoseptica*, expressed in *E. coli*, Sigma Aldrich F8435-50UN) were added to 40 μl of SEC purified YFP-S-RBD (0.24 mg/ml protein concentration), followed by overnight incubation at room temperature. For mass spectrometry, the protein deglycolysation was achieved by incubating 50μl (0.24mg/ml) protein solution with 1 μl of PNgase F (500U, glycerol free) at 37°C overnight.

### Mass spectrometry

LC/MS analysis was performed on a waters LCT Premier mass spectrometer (ESI-TOF) and HPLC Waters 2795. Samples were chromatographed on a Reprosil-PUR 2000 C18-AQ column (3 µm, 100mmX2mm) heated to 50°C using the conditions shown in Table 1:

**Table 1.**
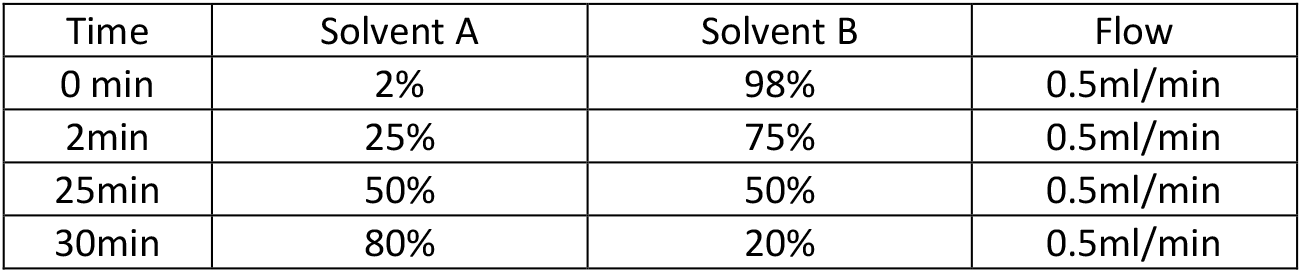
HPLC Method description. Solvent A : acetonitrile with 0.1% formic acid, Solvent B: water with 0.1 % formic acid

| Time | Solvent A | Solvent B | Flow |
| --- | --- | --- | --- |
| 0 min | 2% | 98% | 0.5ml/min |
| 2min | 25% | 75% | 0.5ml/min |
| 25min | 50% | 50% | 0.5ml/min |
| 30min | 80% | 20% | 0.5ml/min |

### SDS-PAGE analysis

NuPAGE 4-12% BisTris, 1.0 mm x 12 well (invitrogen by Thermo Fisher Scientific, NP0322BOX) gels were run in NuPAGE MES SDS running buffer (Invitrogen NP0002) at 150 V. NuPAGE LDS Sample Buffer (Invitrogen) was used with or without DTT. PageRuler Plus Prestained Protein Ladder (Thermo Scientific 26619) was used. Proteins eluted from analytical SEC were concentrated in 10 kDa cutoff spin concentrators prior to analysis by SDS-PAGE. Staining was performed using InstantBlue Coomassie Protein Stain (Expedeon, ISB1L).

## Results and Discussion

### Construct design

Our aim was to produce high-quality, soluble 2019-nCoV spike RBD labeled with a fluorescent protein for easy detection. Spike RBD contains disulfide bonds and N-glycosylations (see e.g. PDB entry 6M17, Yan et al., 2020; PDB entry 6VSB, Wrapp et al., 2020; PDB entry 6LZG, Wang et al. 2020). Therefore, this protein domain is usually produced by secretion from eukaryotic cells.

Only few fusion proteins are commonly used for secreted proteins, notably the constant domain (Fc) of IgG and human serum albumin (Dalton and Barton, 2014). We instead used yellow fluorescent protein as a fusion protein. Analysis of enhanced yellow fluorescent protein (eYFP, Ormö et al., 1996), using the NetNGlyc 1.0 server and the NetOGlyc 4.0 server (Steentoft et al., 2013), revealed no N-glycosylation sites, but a single putative O-glycosylation site, just above threshold, within the YFP sequence. Analysis of the YFP structure showed that the putative O-glycosylation site is near the surface of the protein. Furthermore, secretion of the enhanced green fluorescent protein (eGFP) has previously been described (Román et al., 2016). GFP is nearly identical to YFP in structure and sequence, and also contains the putative O-linked glycosylation site. This same publication (Román et al., 2016) also suggested improved protein secretion levels when using the interferon alpha 2 (IFNα2) signal peptide, compared to a number of commonly used signal peptides, including the signal peptide of interleukin-2 (IL-2). For this reason, we used the IFNα2 signal peptide in our construct, in nearly the same context to the fluorescent protein as in Ref. (Román et al., 2016), except that we left out the start methionine of YFP, since translation starts at the start ATG of the signal peptide upstream of the YFP. Instead, we inserted a short linker (translating into Gly-Ser), which allowed the insertion of a BamHI restriction endonuclease recognition sequence for later use of the vector with the signal peptide for other targets.

The construct was designed for insertion into the HindIII and XbaI sites of the vector pcDNA 4/TO (Invitrogen), a mammalian expression vector that allows tetracycline-inducible expression from a CMV promoter in cells expressing the tetracycline repressor protein, and constitutive expression in cells not containing the tetracycline repressor protein. At the start of the insert, we entered a NotI site containing a partial Kozak sequence (GCGGCCGCC**ATG**G), which we completed with additional nucleotides. The penultimate residue in the signal peptide is alanine, resulting in an optimal **ATG**G DNA sequence (Kozak 1987). A FLAG-tag for detection of the fusion protein or cleaved-off YFP was included at the C-terminus of YFP upstream of a human rhinovirus 3C protease cleavage site. An XhoI restriction endonuclease recognition site was included in the sequence coding for Leu-Glu of the 3C protease site, for later use of the vector with the signal peptide and YFP for other targets. The sequence coding for the 2019-nCoV-spike_RBD with a C-terminal, non-cleavable 8x His-tag and two stop codons, was inserted just downstream of the rhinovirus 3C protease site. The resulting expression construct is depicted in Figure 1.

**Figure 1.**
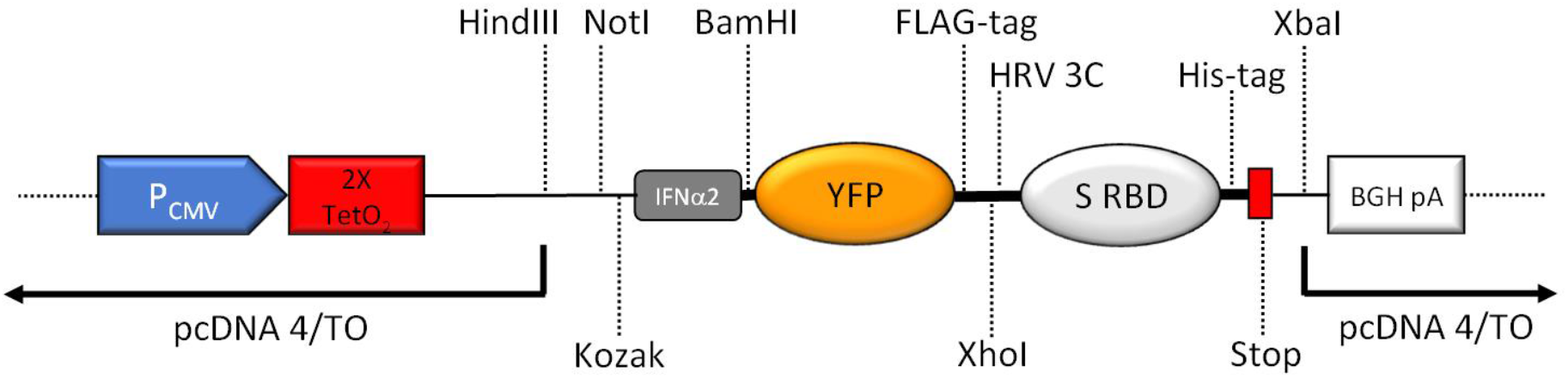
Schematic overview of the key features of the expression region of the construct (not to scale). CMV promoter (P_CMV_), tetracycline operators (2X TetO_2_), Kozak consensus sequence (Kozak), human interferon alpha 2 (IFNα2) signal peptide including start codon for the complete fusion protein, yellow fluorescent protein (YFP), human rhinovirus 3C protease cleavage site (HRV 3C), 2019-nCoV spike RBD (S RBD), 8xHis-tag (His-tag), two stop codons (Stop), BGH polyadenylation sequence (BGA pA). Important unique restriction enzyme recognition sites are indicated.

### Expression

We transfected the expression plasmid into HEK293 cells and generated stable cells by selection with Zeocin. We then expanded the adherent, stably transfected cell culture in a flask with 75 cm^2^ surface area. When a confluence of ∼50% was reached, the DMEM/FBS medium was replaced by serum-free Opti-MEM medium. The supernatant medium was collected three times a week, clarified by centrifugation, supplemented with protease inhibitor, and successively incubated with the same 200 µl Ni-NTA agarose batch. This process was repeated until the Ni-NTA agarose clearly turned yellowish in color. This stage was reached after nine sequential incubations, each with 12-15 ml medium.

We analyzed the YFP fluorescence of each medium batch that we collected. Comparison to the YFP fluorescence of a purified YFP of known concentration allowed an initial estimation of the amount of secreted protein. 12 ml of medium from 48 hours incubation with the confluent culture typically produced a fluorescence peak height of ∼900 relative fluorescence units, which corresponds to a YFP concentration of ∼2 μg/ml.

The cells, originally at ∼50% confluence, reached ∼90-100% confluence within a week in serum-free medium, and a subpopulation of cells detached in confluent cultures and had to be removed from the medium by centrifugation prior to addition to the Ni-NTA resin. After reaching confluence, the amount of protein secreted into the medium remained stable over more than six weeks, based on fluorescence measurements (data not shown).

To upscale protein production, we expanded the stably transfected cells from a backup plate to two larger flasks (150 cm^2^ surface area each). In the original expression flask, the cells had grown slowly after changing to serum-free medium, and the amount of secreted protein increased significantly as the cells reached higher confluence. Furthermore, confluent cultures remained productive for several weeks. For those reasons, we grew the larger scale cultures to confluence before changing to serum-free medium.

We then collected medium from one 75 cm^2^ flask and two 150 cm^2^ flasks and sequentially incubated the collected medium with a 1 ml batch of Ni-NTA agarose until the resin turned yellow (∼280 ml medium total, collected over 14 days).

### Protein purification

The Ni-NTA resin was washed and the protein was eluted in an imidazole-containing buffer. The initial small scale IMAC purification from 200 µl Ni-NTA resin yielded 280 µg protein of high purity (Figure 2 A). The protein solution was monodisperse according to analytical SEC, resulting in a single main SEC peak at the expected retention volume (Figure 2 B).

The upscaled purification from 1 ml Ni-NTA resin yielded 3.3 mg of pure protein after IMAC.

**Figure 2.**
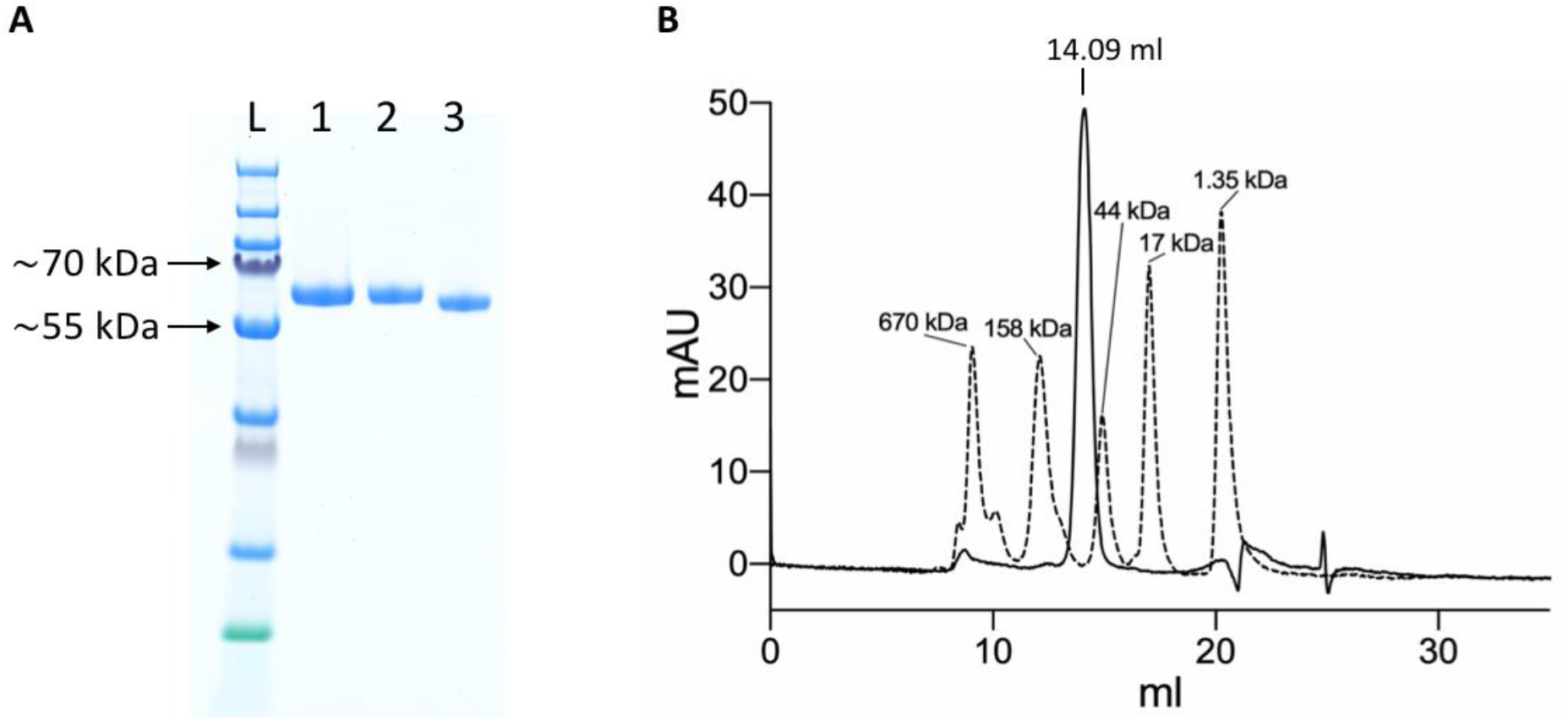
Purification of the YFP-S_RBD fusion protein. A) SDS-PAGE analysis. L, Ladder; 1, YFP-S_RBD from Ni-NTA IMAC; 2, YFP-S_RBD from SEC; 3, YFP-S_RBD from SEC deglycosylated with PNGase F. B) 280 nm absorbance traces from analytical SEC. The trace from a YFP-S_RBD SEC run (solid line) is shown overlaid with the trace of a SEC run with a standard (bio-rad gel filtration standard).

### Protein characterization

Based on analysis by SDS-PAGE, the protein purity was already high after Ni-NTA IMAC. The SEC profile of the YFP-S_RBD fusion protein confirmed the high purity and also showed that the protein solution was monodisperse. This was confirmed by negative staining electron microscopy (EM) (Figure 3 A).

**Figure 3.**
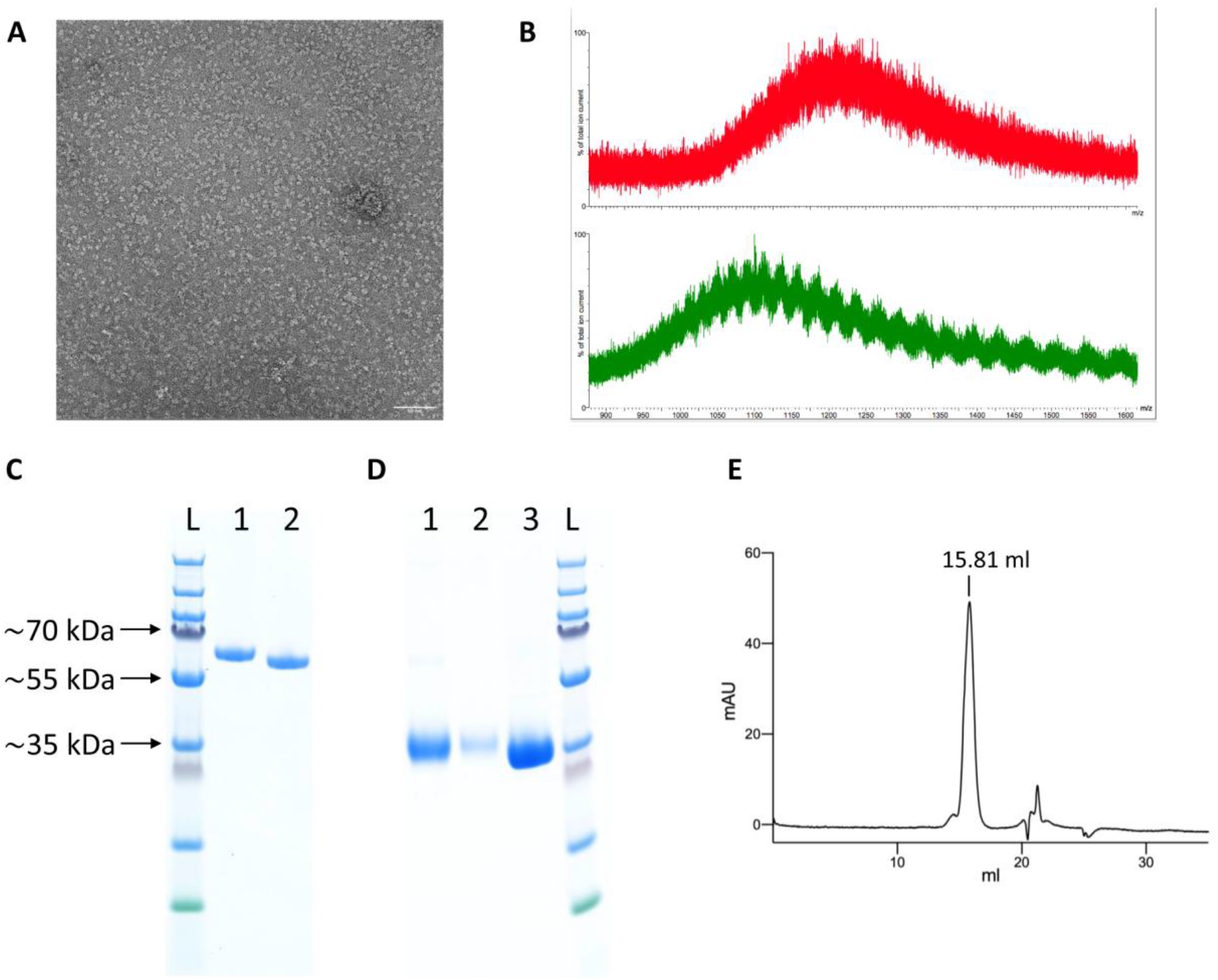
Protein characterization. A) Negative staining electron micrograph of SEC-purified YFP-S_RBD fusion protein. B) Mass spectrometry results of untreated (red, top) and PNGase F treated (green, bottom) YFP-S_RBD fusion protein C, D) SDS-PAGE analysis. C) YFP-S_RBD. L, Ladder; 1, Reducing SDS sample buffer; 2, Non-reducing sample buffer. D) PreScission-cleaved YFP-S_RBD. 1, S_RBD from Ni-NTA IMAC; 2, S_RBD from SEC; 3, YFP washed off the Ni-NTA resin after on-bead cleavage, after removal of the protease. E) Analytical SEC profile (280 nm absorbance) of S_RBD.

Incubation of the purified fusion protein with the enzyme PNGase F resulted in slightly faster migration on SDS-PAGE, confirming the presence of N-glycosylations in the expressed protein (Figure 2 A). Mass spectrometry analysis confirmed the presence of glycosylations and a reduction thereof upon PNGase treatment (Figure 3 B). The fusion protein migrated slightly faster on SDS-PAGE in non-reducing conditions than in reducing conditions, indicating the presence of disulfide bonds in the protein domain (Figure 3 C).

To test whether the S_RBD protein retains its properties after removal of the fluorescent protein tag, the YFP was removed by rhinovirus 3C protease cleavage. 400 μl Ni-NTA resin were loaded with protein in five steps with a total of ∼260 ml expression medium. After washing, the Ni-NTA resin was incubated overnight in the presence of PreScission protease (GST-tagged human rhinovirus 3C protease). The YFP was then washed off and collected, while the His-tagged S_RBD protein remained on the column. The protein was eluted from the now colorless Ni-NTA resin using an imidazole-containing buffer and analyzed by SDS-PAGE (Figure 3 D). The collected cleaved-off YFP was also analyzed on SDS-PAGE, after incubation with Glutathione sepharose 4B to remove the GST-tagged protease (Figure 3 D). The hS_RBD protein was analyzed by analytical SEC. There was a single main peak at the expected retention volume, with only a slight shoulder, confirming that the S_RBD domain retained its solubility and monodispersity after removal of the YFP fusion protein (Figure 3 E).

To test whether the purified YFP-S_RBD fusion protein binds its target receptor ACE2, we produced and purified human ACE2 peptidase domain and analyzed the separate proteins as well as the complex of the two proteins by analytical SEC experiments. The complex co-eluted in a peak at a reduced retention volume compared to the peak from ACE2 run alone or the peak of YFP-S_RDB run alone, clearly confirming complex formation (Figure 4 A-C). To test whether the S_RBD domain retains its ACE2-binding activity after proteolytic removal of the YFP, the analytical SEC experiment was repeated with PreScission protease cleaved, purified S_RBD. The two proteins co-eluted in a peak at a reduced retention volume compared to the separate proteins, confirming binding (Figure 4 D and E).

**Figure 4.**
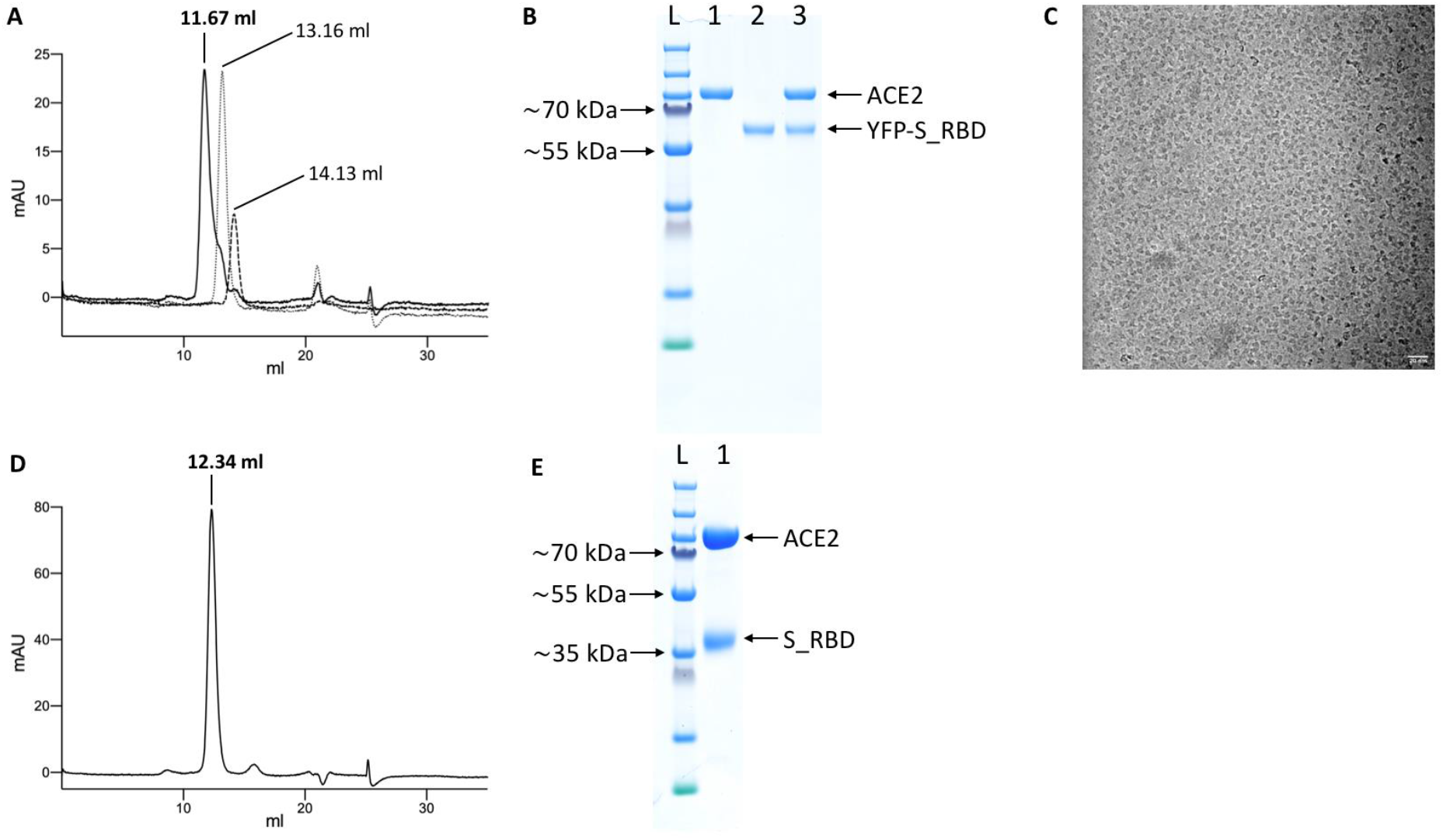
Analysis of complex formation by analytical SEC. A) Overlay of the 280 nm absorbance SEC traces from the complex (YFP-S_RBD / ACE2 run together, solid black line), ACE2 run separately (fine dotted line) and YFP-hS_RBD run separately (coarse dotted line). B) SDS-PAGE analysis of the proteins. L, Ladder; 1, ACE2; 2; YFP-S_RBD; 3, complex eluted from SEC. C) Cryo-EM micrograph of the ACE2/YFP-S_RBD complex. D) 280 nm absorbance SEC trace of the complex of ACE2 with S_RBD (after proteolytic removal of YFP). E) SDS-PAGE analysis of the complex eluted from (D).

## Author contributions

The project was initiated and coordinated by R.M.B.. The ACE2 construct was sub-cloned by T.B.. Cell culture experiments, protein purification and biochemical analysis were performed by R.M.B., T.B. and G.C.. Electron microscopy experiments were carried out by E.P. and T.B.. Mass spectrometry analysis was performed by A.B.. The manuscript was written by R.M.B. and T.B. with contributions from all authors.

## Acknowledgements

We thank Takashi Ishikawa, Gebhard F.X. Schertler and Michel Steinmetz for supporting the project. This work was in part supported by grants from UBS Promedica (1401/M) and the Swiss National Science Foundation (SNF SPARK, CRSK-3_190414) to R.M.B., and by a Sinergia grant from the Swiss National Science Foundation (CRSII5_183563 / 1). We furthermore acknowledge support in part from the PSI COVID19 Emergency Science Fund.

## References

Dalton AC, Barton WA. Over-expression of secreted proteins from mammalian cell lines. Protein Sci. 2014 May;23(5):517–25. doi: 10.1002/pro.2439. Epub 2014 Mar 11. PMID: 24510886; PMCID: PMC4005704.

Kozak M. An analysis of 5’-noncoding sequences from 699 vertebrate messenger RNAs. Nucleic Acids Res. 1987 Oct 26;15(20):8125–48. doi: 10.1093/nar/15.20.8125. PMID: 3313277; PMCID: PMC306349.

Li F. Structure, Function, and Evolution of Coronavirus Spike Proteins. Annu Rev Virol. 2016 Sep 29;3(1):237–261. doi: 10.1146/annurev-virology-110615-042301. Epub 2016 Aug 25. PMID: 27578435; PMCID: PMC5457962.

Ormö M, Cubitt AB, Kallio K, Gross LA, Tsien RY, Remington SJ. Crystal structure of the Aequorea victoria green fluorescent protein. Science. 1996 Sep 6;273(5280):1392–5. doi: 10.1126/science.273.5280.1392. PMID: 8703075.

Román R, Miret J, Scalia F, Casablancas A, Lecina M, Cairó JJ. Enhancing heterologous protein expression and secretion in HEK293 cells by means of combination of CMV promoter and IFNα2 signal peptide. J Biotechnol. 2016 Dec 10;239:57–60. doi: 10.1016/j.jbiotec.2016.10.005. Epub 2016 Oct 7. PMID: 27725209.

Steentoft C, Vakhrushev SY, Joshi HJ, Kong Y, Vester-Christensen MB, Schjoldager KT, Lavrsen K, Dabelsteen S, Pedersen NB, Marcos-Silva L, Gupta R, Bennett EP, Mandel U, Brunak S, Wandall HH, Levery SB, Clausen H. Precision mapping of the human O-GalNAc glycoproteome through SimpleCell technology. EMBO J. 2013 May 15;32(10):1478–88. doi: 10.1038/emboj.2013.79. Epub 2013 Apr 12. PMID: 23584533; PMCID: PMC3655468.

Toelzer C, Gupta K, Yadav SKN, Borucu U, Davidson AD, Kavanagh Williamson M, Shoemark DK, Garzoni F, Staufer O, Milligan R, Capin J, Mulholland AJ, Spatz J, Fitzgerald D, Berger I, Schaffitzel C. Free fatty acid binding pocket in the locked structure of SARS-CoV-2 spike protein. Science. 2020 Sep 21:eabd3255. doi: 10.1126/science.abd3255. Epub ahead of print. PMID: 32958580.

Wang Q, Zhang Y, Wu L, Niu S, Song C, Zhang Z, Lu G, Qiao C, Hu Y, Yuen KY, Wang Q, Zhou H, Yan J, Qi J. Structural and Functional Basis of SARS-CoV-2 Entry by Using Human ACE2. Cell. 2020 May 14;181(4):894-904.e9. doi: 10.1016/j.cell.2020.03.045. Epub 2020 Apr 9. PMID: 32275855; PMCID: PMC7144619.

Wrapp D, Wang N, Corbett KS, Goldsmith JA, Hsieh CL, Abiona O, Graham BS, McLellan JS. Cryo-EM structure of the 2019-nCoV spike in the prefusion conformation. Science. 2020 Mar 13;367(6483):1260–1263. doi: 10.1126/science.abb2507. Epub 2020 Feb 19. PMID: 32075877; PMCID: PMC7164637.

Yan R, Zhang Y, Li Y, Xia L, Guo Y, Zhou Q. Structural basis for the recognition of SARS-CoV-2 by full-length human ACE2. Science. 2020 Mar 27;367(6485):1444–1448. doi: 10.1126/science.abb2762. Epub 2020 Mar 4. PMID: 32132184; PMCID: PMC7164635.

